# *MetaNovo:* a probabilistic approach to peptide discovery in complex metaproteomic datasets

**DOI:** 10.1101/605550

**Authors:** Matthys G Potgieter, Andrew JM Nel, Suereta Fortuin, Shaun Garnett, Jerome M. Wendoh, David L. Tabb, Nicola J Mulder, Jonathan M Blackburn

## Abstract

**Background:** Microbiome research is providing important new insights into the metabolic interactions of complex microbial ecosystems involved in fields as diverse as the pathogenesis of human diseases, agriculture and climate change. Poor correlations typically observed between RNA and protein expression datasets make it hard to accurately infer microbial protein synthesis from metagenomic data. Additionally, mass spectrometry-based metaproteomic analyses typically rely on focussed search libraries based on prior knowledge for protein identification that may not represent all the proteins present in a set of samples. Metagenomic 16S rRNA sequencing will only target the bacterial component, while whole genome sequencing is at best an indirect measure of expressed proteomes. We describe a novel approach, **MetaNovo**, that combines existing open-source software tools to perform scalable *de novo* sequence tag matching with a novel algorithm for probabilistic optimization of the entire **UniProt** knowledgebase to create tailored databases for target-decoy searches directly at the proteome level, enabling analyses without prior expectation of sample composition or metagenomic data generation, and compatible with standard downstream analysis pipelines.

**Results:** We compared **MetaNovo** to published results from the **MetaPro-IQ** pipeline on 8 human mucosal-luminal interface samples, with comparable numbers of peptide and protein identifications, many shared peptide sequences and a similar bacterial taxonomic distribution compared to that found using a matched metagenome database - but simultaneously identified many more non-bacterial peptides than the previous approaches**. MetaNovo** was also benchmarked on samples of known microbial composition against matched metagenomic and whole genomic database workflows, yielding many more MS/MS identifications for the expected taxa, with improved taxonomic representation, while also highlighting previously described genome sequencing quality concerns for one of the organisms, and identifying a known sample contaminant without prior expectation.

**Conclusions:** By estimating taxonomic and peptide level information directly on microbiome samples from tandem mass spectrometry data, **MetaNovo** enables the simultaneous identification of peptides from all domains of life in metaproteome samples, bypassing the need for curated sequence search databases. We show that the **MetaNovo** approach to mass spectrometry metaproteomics is more accurate than current gold standard approaches of tailored or matched genomic database searches, can identify sample contaminants without prior expectation and yields insights into previously unidentified metaproteomic signals, building on the potential for complex mass spectrometry metaproteomic data to speak for itself. The pipeline source code is available on **GitHub**^1^ and documentation is provided to run the software as a singularity-compatible docker image available from the **Docker Hub**^2^.

## Introduction

Characterising microbial ecosystems from clinical or environmental samples promises insights into the complex metabolic pathways involved in processes as diverse as carbon sequestration and climate change to the susceptibility and progression of human diseases [1]. Microbial communities play a wide-ranging role in the complex determinants of human well-being, with potential disease implications should the balance be disrupted [2].

Genome and transcriptome sequencing approaches to microbiome analysis allow for functional and taxonomic characterization of the genes and organisms involved in complex microbial communities, but it has been shown that measures of gene transcription do not generally correlate well with measured protein abundance [3]. On the other hand, metaproteomic and clinical proteomic approaches in principle allow researchers to obtain a snapshot of all the proteins present in a complex microbiome sample at a given time, providing a quantitative window into key functional components of complex pathways and organism interactions, including the direct measurement of dynamic changes in microbial protein composition, localisation and modification that may mediate host/pathogen interactions in the context of human health and disease.

Classically, the identification of peptides (and by inference, the parent proteins) from complex mass spectrometry datasets has been largely dependent on the availability of focussed, representative and relatively small sequence databases or spectral libraries against which to search tandem mass spectra - perhaps due in part to the historical emphasis in the field on the analysis of model organism proteomes to answer simple biological questions. However, the analysis of complex, multi-species samples – such as human microbiome samples where the total number of microbial genes may vastly exceed the number of human genes - is far less straightforward for a variety of reasons, including: the vast majority of specific organisms in any given microbiome are likely to not have been cultured, identified, or characterised in the laboratory, so appropriate reference genomes may not exist to underpin proteomic data analyses; and the fact that massive expansion of proteomic sequence databases used in target-decoy-based peptide-spectrum matching has been shown to lead to dramatically reduced identification rates due to higher rates of false negatives and false discovery misestimation problems [4].

Various methods have been developed to address the challenges of metaproteomic MS/MS identification. Previous methods for metaproteomic analysis have addressed these issues essentially by relying on prior expectation of the microbial composition of a given microbiome sample, for example using 16S sequencing data to produce a focussed metaproteome library to search [5]. However, some limitations of this approach are that: (i) it assumes that the 16S data itself provides a complete description of all bacteria present in the sample, which is probably not true; (ii) it assumes that appropriate reference bacterial proteomes exist and are close enough in sequence to the clinical isolate proteomes for proteomic data analysis to be valid, which may also not be true; (iii) it automatically ignores all non-bacterial components of the microbiome; & (iv) the expressed proteome even within a single species is dynamic and not constant under all conditions, leading to the likely inclusion of spurious protein sequences even for correctly included organisms, thus inflating the search space and leading to decreased sensitivity. These approaches also imply additional analyses that depend on resources and samples that may not be available in all cases. Whole metagenome sequencing itself also suffers from the complexity of short reads leading to fragmented contigs and scaffolds which may interrupt protein-coding regions and hamper CDS predictions. Tang et al. [6] used *de Bruijn* graphs generated by metagenome assembly to produce protein databases for metaproteomics, where identified peptides may span separate contigs in the graph, yielding many more peptide identifications than linear approaches to metagenome data.

Another powerful method of increasing sensitivity of metaproteome database searches are iterative searches such as the “two-step” method, where matches from a primary search against a very large database are used to create a focused database for target-decoy search yielding a higher number of identifications [7]. Zhang et al [8] developed the **MetaPro-IQ** pipeline for iterative database searching, using a relaxed FDR for the first search to narrow down the database for an FDR-controlled second search. Using this approach to search canonical catalogues of genes to create targeted databases for peptide spectral matching allows researchers to leverage large amounts of publicly available and curated sequence data yielding comparable results to matched metagenomic approaches, however, identifications from curated gene lists may not accurately reflect sample-specific clinical polymorphisms at the peptide sequence level (leading to missed assignments) or even exclude unexpected or uncharacterized organisms entirely, leading to decreased comprehensiveness of the database search space with a corresponding increase in the risk of mis assignments and loss of identification sensitivity. Furthermore, correctly identified peptides may belong to multiple homologous proteins, further complicating protein inference and quantification [9]. These approaches typically rely on conventional peptide spectral matching tools that are not designed for extremely large database sizes such as **UniProtKB** due to the computational resources required, making iterative searches of such large databases prohibitive or too time-consuming, and therefore still rely on some level of filtering or prior knowledge for the first search.

Particular challenges faced by adult gut metaproteomics include inter-subject variability [10], combined with the need to factor in the influence of a very diverse and variable human diet that directly contributes to the protein component of the gut. Thus, whilst matched metagenome bacterial sequencing is the current gold standard to create sequence databases to search metaproteomic mass spectrometry data against, such data is not always available nor necessarily complete. Moreover, such curated databases for a given microbiome typically do not include viral, fungal, host or other eukaryote proteomes that may be present, nor do they account for possible sequence polymorphisms compared to reference strains.

Further pitfalls of the use of targeted databases for metaproteomics include the danger of spurious assignments when proteins present in the data are not included in the search database. A case in point are spectra assigned to viral proteins identified and published in a study but later identified as *Apis mellifera* honey bee proteins by another group - that may have been differently assigned if the original authors included those proteins in the original search database [11]. This scenario underlines a dangerous and seemingly ignored fact in many published metaproteomics studies that make no mention of whole domains of life that are known to be present in samples being analysed, such as viruses, archaea, fungi and dietary components in gut metaproteomics analyses that merely use a database targeting bacterial proteins. This may have a knock-on effect on downstream research that rely on the findings. The risk of unique and rare contaminants from environmental or clinical samples being misassigned due to a non-representative list of putative contaminants, further complicates the situation, as prior expectation of sample composition cannot be expected to cater for these.

The ideal database for mass spectrometry-based peptide identification in metaproteomic datasets would therefore be comprehensive, whilst excluding proteins that are genuinely absent, and would not be reliant on prior expectation of sample composition, nor would it require prior generation of DNA/RNA sequencing data to inform the construction of a search database. High quality curated data repositories like **SwissProt** are arguably also biassed at the level of database composition towards culturable and model organisms that are frequently studied, and can not be expected to represent all the sequence diversity of clinical isolates. Large protein databases like **NCBI nr** and **trEMBL** include automatically generated annotations from genomics data, and are more comprehensive but arguably also biased by factors including biases in the gene prediction algorithms themselves, and these need to be factored into downstream approaches.

Taxonomic profiling of proteins identified during metaproteomic analysis is complicated by high levels of shared tryptic peptides between homologous proteins of closely related organisms. The UniPept *pept2lca* algorithm [12] assigns the lowest common ancestor (LCA) for a given peptide assignment. The recently published **ProteoClade** algorithm [13] allows users to apply species level annotation to identified peptides using customised search databases.

*De novo* sequencing of peptides from mass spectrometry data has long been used for database filtration, allowing for rapid searches of very large search spaces with sequence tags prior to peptide-spectral matching [14]. Tanner et al. published the **InsPecT** tool in 2005 [15]. These approaches work by reducing the database size by orders of magnitude, while retaining the correct sequences with high probability [15]. The sequence tag filtering approach is interesting, as although in isolation full peptide sequencing of Higher-energy C-trap dissociation (HCD) data has been found to be around 35% accurate [16], when a database match does occur the validity of those sequences become more likely. Further, as only a few correct proteotypic peptides are needed to confidently identify a protein [16], and thus select it for inclusion in the filtered database, high sequence coverage is less important at this step. As incomplete fragmentation of experimental spectra hampers accurate full-length sequencing using *de novo* algorithms [17], partial sequencing with sequence tag approaches with tools such as **DirecTag** [18] are a robust and scalable approach for database filtering. Recent advances in deep-learning-based approaches have significantly improved the accuracy of full sequence *de novo* sequencing approaches, with tools such as **pNovo3** yielding up to 89% improvement in precision when compared to other state-of-the-art tools, illustrating the current rapid progress in this field [19]. Particularly considering significant inter-subject variation and the polymorphism of clinical strains, *de novo* sequencing has been suggested as a viable strategy for peptide identification in gut metaproteomics [20]. However, scalable and robust *de novo* sequencing-based pipelines need to be able to process the rapidly expanding amount of proteomics and genomics information available. Further, protein inference based on identified peptides from the above approaches requires scalable and accurate approaches for metaproteomics.

Here we present a novel metaproteomic analysis pipeline - **MetaNovo** - extending the conventional methodology for iterative database search by mapping *de novo* sequence tags to very large protein sequence databases using a high-performance and parallelized computing pipeline as the first search, generating compact databases for target-decoy analysis and False Discovery Rate (FDR) controlled protein identification suitable for integration with existing MS/MS analysis pipelines. We compare the results of **MetaNovo** against those published for the two-step **MetaPro-IQ** pipeline [8] using a publicly available mass spectrometry dataset.

We further validated the pipeline on samples of known microbial composition as a ground truth. To our knowledge, this is the first published workflow for iterative two-step searching that uses *de novo* sequencing and probabilistic protein inference as a first step, followed by a conventional target-decoy search with FDR control in the second. We show that replacing conventional database searching with *de novo* sequencing in the first step extends the applicability of iterative searches to extremely large databases, which have to date only been applied to large curated lists of proteins, thus decreasing the bias inherent in the analysis of complex metaproteomic data using these approaches.

## Design and Implementation

### ***MetaNovo*** database generation workflow and benchmarking

#### UniProt database statistics

The 2019_11 release [21] of **UniProt** was downloaded, and a combined FASTA database created from **UniProt SwissProt**, **TREMBL**, and **SwissProt** alternative splice variant sequences (ca. 180 million sequences). See *Supplementary Table 1. Taxonomic distribution by Kingdom of the **UniProt** database*.

**General software overview.** The **MetaNovo** software is a configurable command-line linux **Bash** pipeline that combines open-source software tools with custom **Python** libraries and scripts to provide targeted protein identification search libraries in **FASTA** format. The pipeline has three main components based on custom and existing open-source tools - generating *de novo* sequence tags using **DirecTag**, mapping the tags to a protein sequence database using **PeptideMapper**, and a novel algorithm for probabilistic protein ranking and filtering based on estimated species and protein abundance. The **MetaNovo** software is available from **GitHub** [22] and can be run as a standalone **Singularity** or **Docker** container available from the **Docker Hub** [23]. All results have been uploaded to the **ProteomeXchange Consortium** via the **PRIDE** [24] partner repository with the dataset identifier **PXD030708**. *See Figure 1. The Typical **MetaNovo** workflow*.

**Figure 1.**
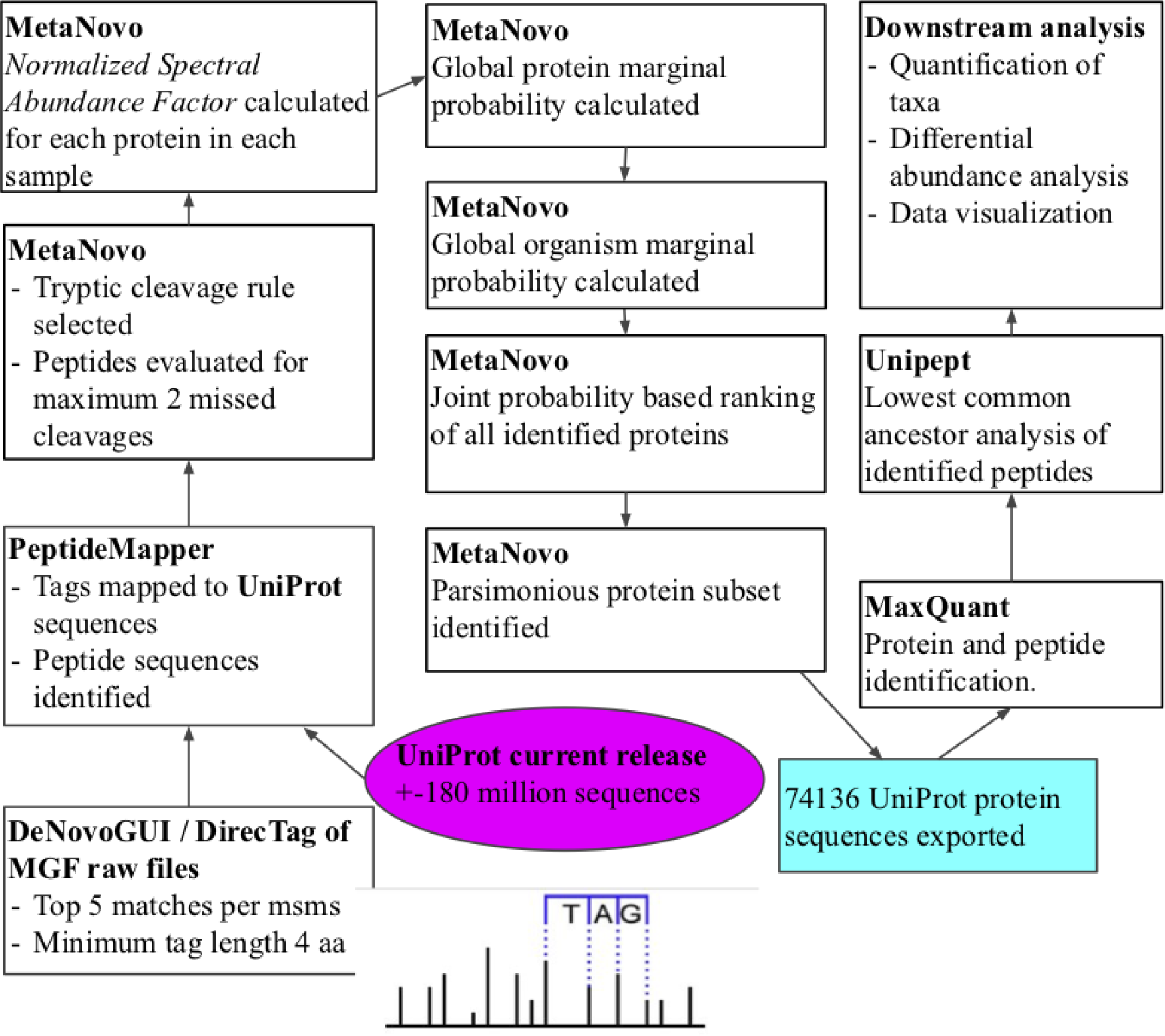
Visualisation of the **MetaNovo** workflow used to analyse the mass spectrometry data of 8 human mucosal-luminal interface samples. Raw mass-spectrometry data were analysed using the **MetaNovo** pipeline in MGF format, using *de novo* sequence tags to create a targeted FASTA file for target-decoy search.

**Figure 2.**
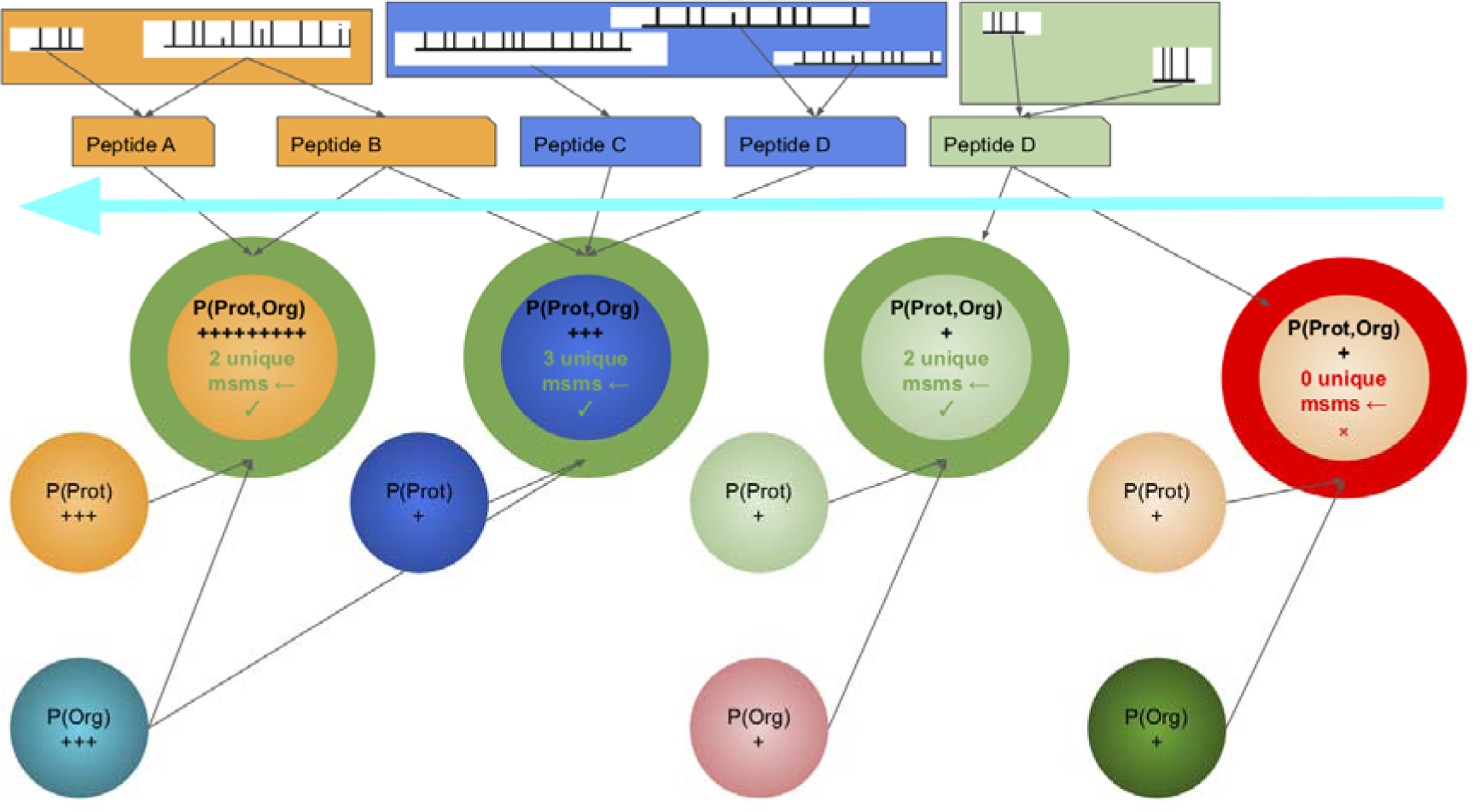
A Graphical representation of the **MetaNovo** algorithm applied for database filtration. *Normalised spectral abundance factor* calculations include non-unique spectra. Probabilities are represented by +’s. The number of unique spectra for each protein is determined based on its position in the ranked list, and only include spectra that do not appear in the set of proteins in the list above (but may include spectra that appear below), such as **Peptide B** that is counted towards the first protein in the list, but not the second. Tiebreaks for adjacent and nearly identical isoforms that share the same set of spectra, will be based on the shortest (most probable) sequence having a higher NSAF (and thus a higher protein probability). Proteins in green will be selected for inclusion in the filtered database, and proteins in red will be excluded (no unique spectra).

#### Generation of *de novo* sequence tags

Raw MS/MS files need to be converted to *Mascot Generic Feature* (MGF) format prior to analysis. *De novo* sequence tags are generated using **DeNovoGUI** version 1.15.11 [25], with **DirecTag** [18] selected as the sequencing engine. General **MetaNovo** parameters selected for the analyses include fragment and precursor ion mass tolerance of 0.02 Da, with fixed modification ‘*Carbamidomethylation of C’* and variable modifications ‘*Oxidation of M’* and *‘Acetylation of protein N-term’*. For **DirecTag**, a tag sequence length of 4 amino acids was required, and the top 5 sequence tags per spectrum were selected (taking into account alternative possible charge states). These settings were chosen as preliminary runs with a sequence tag length of 3 proved impractical with very large databases, requiring more than 5 days with 24 cores on a high-performance cluster to search 8 MGF files against **UniProt**. Similarly, increasing sequence tag options per spectrum from 5 to 30 considerably increased processing time without a corresponding gain in sensitivity, possibly due to the inclusion of low-scoring matches. The output of **DirecTag** is parsed with a custom **Python** script and all sequence tags for each MS/MS across replicates are stored in an **SQLite3** database. The distinct set of sequence tags (N-terminus mass gap, amino acid sequence, and C-terminus mass gap) are obtained using an **SQL** query, combining identical tags across multiple MS/MS into a single non-redundant list.

#### Mapping sequence tags to a FASTA database

**PeptideMapper** [26] included in **compomics-utilities** version 4.11.19 [27] is used to search the sequence tag set against the protein sequence database. The same mass error tolerance and post-translational modification settings are used as for **DeNovoGUI**, and specified in the config file. The open-source **GNU parallel** tool [28] is used to search the set of sequence tags against the FASTA database in configurable chunk sizes, using a configurable number of cores per node in parallel.

#### Enzymatic cleavage rule evaluation

A custom **Python** script is used to evaluate the cleavage rule of the peptide sequences of identified sequence tags. Only peptides passing the selected cleavage rule are selected for downstream analysis (‘Trypsin, no P rule’ was selected for the benchmarking analysis - cleavage after all *Lys* or *Arg*) with up to 2 missed cleavages allowed. The corresponding sequence tags for the validated peptide sequences are queried against the **SQLite3** [29] database to obtain a mapping of MS/MS ids to protein ids, allowing for the estimation of protein abundance based on mapped MS/MS in a similar manner to spectral counting in a target-decoy search.

#### Normalised spectral abundance factor calculation

Protein abundance is estimated using the *Normalised Spectral Abundance Factor* (NSAF) approach developed for shotgun proteomics [30]. The set of MS/MS IDs in each sample per protein is obtained, and the number of mapped MS/MS ids is divided by protein length to obtain a *Spectral Abundance Factor* (SAF). The SAF values for each sample are divided by the sum of the SAF values in that sample, to obtain an NSAF for each protein in each sample/replicate.

#### Probabilistic ranking of database proteins

The **MetaNovo** algorithm uses the concept of unconditional or marginal probability, which is a probabilistic value generated for each protein in the **UniProt** database corresponding to the probability that a given peptide fragment sampled at random from the raw data will belong to that protein given the available data. To calculate the marginal probabilities, protein NSAF values are summed across replicates, and divided by the sum of all NSAF values across all replicates, with a maximum value of 1 for each protein. This value is equivalent to the proportion that a given protein makes up of all replicates by summed NSAF value. Database proteins are ranked by this marginal probability and filtered such that a minimal set of ranked proteins is obtained where each protein in the list contains at least one unique MS/MS scan ID relative to the set of proteins above its position in the list - and excluding all proteins that do not contain any uniquely identified spectra relative to the set of proteins above their position in the ranked list. Using this methodology, sets of protein isoforms that share an exact set of mapped MS/MS ids will be represented by the shortest sequence in the set due to having a higher NSAF value, based on the principle of Occam’s razor where longer sequences are more likely to have sequence tag matches by pure chance. The organism names are obtained from the **UniProt** FASTA headers, and summed NSAF values in the filtered set are aggregated by organism and divided by the total to obtain an organism level marginal probability. Following this, the proteins in the unfiltered set are annotated with their respective calculated organism marginal probability. The unfiltered set of proteins is re-ranked based on the joint probability of the organism and protein probability values, based on the simplistic assumption of conditional independence (Naive Bayes Assumption) which allows low abundance proteins with a high estimated relative abundance at the organism level to be favoured compared to proteins where both the protein and organism abundance are estimated to be low. Benefits of the Naive Bayes assumption of conditional independence between predictors are ease of mathematical implementation, and being highly scalable to the number of predictors and data points, without sacrificing model accuracy, and allowing model training and classification with a single pass over the data.

Following the second ranking step, the database is re-filtered to obtain a minimal subset of proteins that can explain all mapped *de novo* sequence tags, based on estimated relative protein and organism abundance.

### Mucosal-luminal interface samples

#### Mucosal-luminal interface (MLI) sample metagenomic and proteomics data

Metagenome and proteomics data of 8 MLI samples from adolescent volunteers obtained during colonoscopy were downloaded from **PRIDE** with identifier **PXD003528** and through author correspondence. The sample processing, mass spectrometry and metagenomics database creation methods have already been described [8].

#### *MetaNovo* database generation

The December 2019 release [21] of **UniProt** (containing ca. 180 million protein entries) was used to create a database containing 74136 entries. The **MetaNovo** settings described above were used.

#### Database search using MaxQuant

**MaxQuant** version 1.5.2.8 was used to search the **MetaNovo** database, with the same settings as for the **MetaPro-IQ** publication. *Acetyl (Protein N-term)* and *Oxidation (M)* were selected as variable modifications, and *Carbamidomethyl (C)* was selected as a fixed modification. Specific enzyme mode with *Trypsin/P* was selected with up to 2 missed cleavages allowed, and a PSM and protein FDR of 0.01 was required.

### 9MM Samples

#### 9MM sample and validation databases

Two samples of known microbial mixture from a single biological replicate (9MM) were obtained from **PeptideAtlas** at identifier **PASS00194**. Detailed proteomic methods are described in the original publication [31]. In brief, each of 9 organisms were cultured separately on the appropriate growing media, and divided into aliquots of approximately 10 CFU each. The 9MM sample was created by combining an aliquot of each organism pellet, and processed separately by filter-aided sample preparation (FASP) and protein precipitation followed by in-solution digestion (PPID). Two databases were selected from the original publication for comparison. One, the top-performing database from the previous publication was created by single genome assembly followed by gene prediction and protein annotation using **TrEMBL (SGA-PA**). Secondly, to illustrate a typical metaproteomics workflow, the database created by NGS of the combined sample (metagenome sequencing) followed by gene prediction and protein annotation using **TrEMBL** was selected (**Meta-PA**).

#### *MetaNovo* database generation

The December 2019 release [21] of **UniProt** (containing ca. 180 million protein entries) was used to create a database containing 13195 entries. The **MetaNovo** settings described above were used.

**Database search using MaxQuant. MaxQuant** version 1.5.2.8 was used to search the **MetaNovo, SGA-PA** and **Meta-PA** databases using the same search parameters. *Acetyl (Protein N-term)* and *Oxidation (M)* were selected as variable modifications, and *Carbamidomethyl (C)* was selected as a fixed modification. Specific enzyme mode with *Trypsin/P* was selected with up to 2 missed cleavages allowed, and a PSM and protein FDR of 0.01 was required. The FASP and PPID samples were treated as separate experiments.

### Bioinformatic analysis

#### Posterior error probability (PEP) score analysis

PEP scores are a standard metric to identify and rank the quality of PSMs and are used during the FDR calculation of target decoy searches, as false-positive PSMs tend to rank similarly to reverse or decoy hits, allowing a convenient method to estimate the FDR at a given position in the PEP score ranking [32]. PEP scores have been applied in the field of proteogenomics to evaluate the accuracy of novel peptide identifications, by comparing their distribution to that of reverse hits, allowing for the assessment of the level completeness of protein annotation in a given strain, as novel peptide identifications in completely annotated genomes are likely false positive identifications [33]. The PEP scores of PSMs obtained using **MaxQuant** were obtained from the *peptides.txt* output files of the different database runs of the MLI samples - using **MetaNovo**, **MetaPro-IQ** using the integrated gene catalog (IGC) of human gut microbial genes [34], hereafter referred to as **MetaPro-IQ/IGC**, and a matched metagenome database hereafter referred to as **MetaPro-IQ/Metagenome** (the published **MaxQuant** results using the **MetaPro-IQ/IGC** and **MetaPro-IQ/Metagenome** databases were obtained from PRIDE). PSMs for each run were grouped into “exclusive” peptide sequences only identified in that run, and PSMs of “shared” peptides that were also identified in the other two runs, as well as the reverse hits for each group. Significant differences between the groups were tested for using the *Kruskal-Wallis* non-parametric analysis of variance test followed by *Dunn’s* post hoc test using a custom python script. PEP score distributions were visualised using the python *matplotlib* library.

#### Taxonomic analysis

Peptide sequences were assigned to the lowest common ancestor (taxonomic level) using the **UniPept *pept2lca*** tool, to allow for a comparison between the results of the different approaches and to the ground truth samples of known taxonomic composition (9MM) [31]. Peptide and spectral counts were aggregated at selected phylum levels reported in the ***pept2lca*** output, with custom visualisations done using the Python *matplotlib* module. We followed a previously described approach to filtering spurious taxa based on **UniPept** results by only including taxa that represent more than 0.5 % of taxonomically characterised peptides [31], applied separately at each taxonomic level.

#### BLAST analysis of selected peptides and proteins from 9MM samples

Peptides identified in the 9MM analysis that were assigned using the **UniPept** *pept2lca* tool to organisms not expected from the known mixture were selectively subjected to **NCBI BLAST** analysis. Leading proteins for the same peptides were extracted from the *proteinGroups.txt* file and the protein sequences submitted for **NCBI BLAST** [35]. Proteotypic peptides were defined as only matching to a single species with 100 % identity in the **NCBI** nr database. Species-specific protein sequences were defined as those sequences that only matched to a single species above a threshold of 90 % identity in the **NCBI** nr database.

#### 9MM Taxonomic representation error scores calculation

Spectral counts obtained from the **MaxQuant** output files were aggregated to *family*, *genus* and *species* levels for the expected taxa. For each expected taxon, the percentage relative to the total characterised MS/MS (including mis assignments) was calculated for the **MetaNovo**, **SGA-PA**, and **Meta-PA** workflows respectively. As approximately equal proportions of the 9 organisms were analysed based on CFU counts, ground truth percentages were assigned to the 3 taxonomic levels accordingly. Mean Squared Error (MSE) scores were calculated for relative taxon abundance for each of the three runs relative to the ground truth percentages and compared. MSE values closer to 0 indicate lower error rates, and therefore a more accurate representation of relative taxon abundance in the samples.

## Results

### MetaNovo identifies non-bacterial taxa, whilst providing comparable bacterial composition data to a matched metagenome approach in human Mucosal-luminal interface samples

#### Protein and peptide identifications

The **MetaNovo** pipeline was run using the entire, unfiltered **UniProt** database (December 2019 release; ca. 180 million entries), resulting in the identification of 69878 target peptides and 15731 protein groups, with 36.37 % of total MS/MS identified. These results are comparable but slightly better with >2% more spectra assigned than those of previous approaches, with 34 and 33% MS/MS identification rates reported by the **MetaPro-IQ** authors using a matched metagenome and the integrated gene catalog (IGC), respectively. 48705 peptides were identified in common by all three runs, while 4873, 6525 and 14049 were identified exclusively by the **MetaPro-IQ/IGC**, **MetaPro-IQ/Metagenome**, and **MetaNovo** databases, respectively. *See Figure 4 B*.

**Figure 4.**
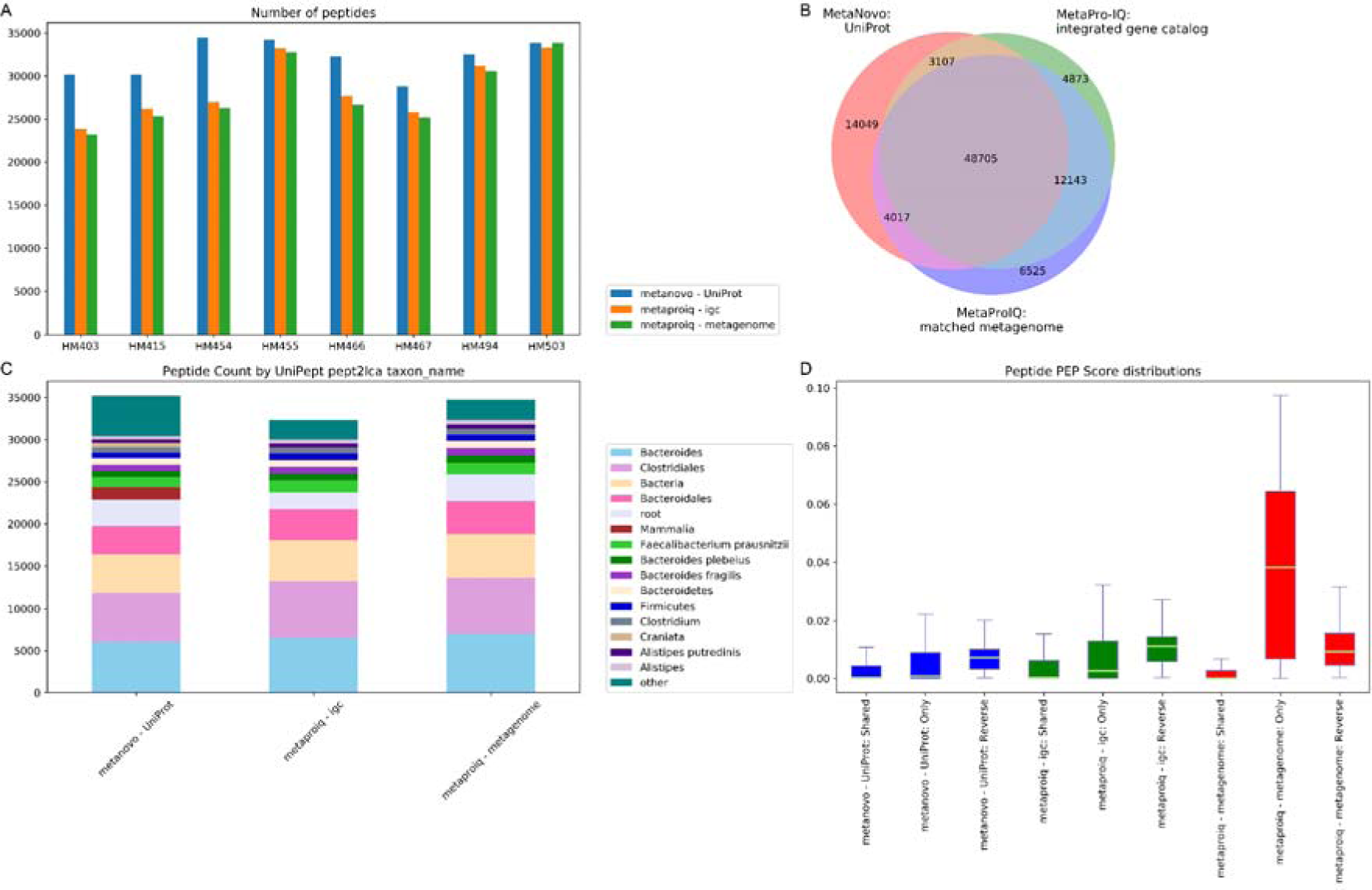
**A.** Bar chart of peptide identifications. The identification rates of **MetaNovo** are comparable to the previously published results of **MetaPro-IQ** using matched metagenome and *integrated gene catalog* databases. **C.** Venn diagram showing large overlap in identified sequences using different approaches, with the highest number of sequences identified using **MetaNovo. C.** Peptide counts by *UniPept lowest common ancestor* showed similar taxonomic distributions obtained from different approaches. **D.** Peptides uniquely identified by MaxQuant using the **MetaNovo** database had a significantly different distribution compared to reverse hits (p-value 6.33e-26), indicating true positive identifications not found using the other two approaches. The boxes extend from the lower to the upper quartile, and the whiskers represent 1.5 times the interquartile range (IQR) below and above the first and third quartiles, respectively.

**Figure 5.**
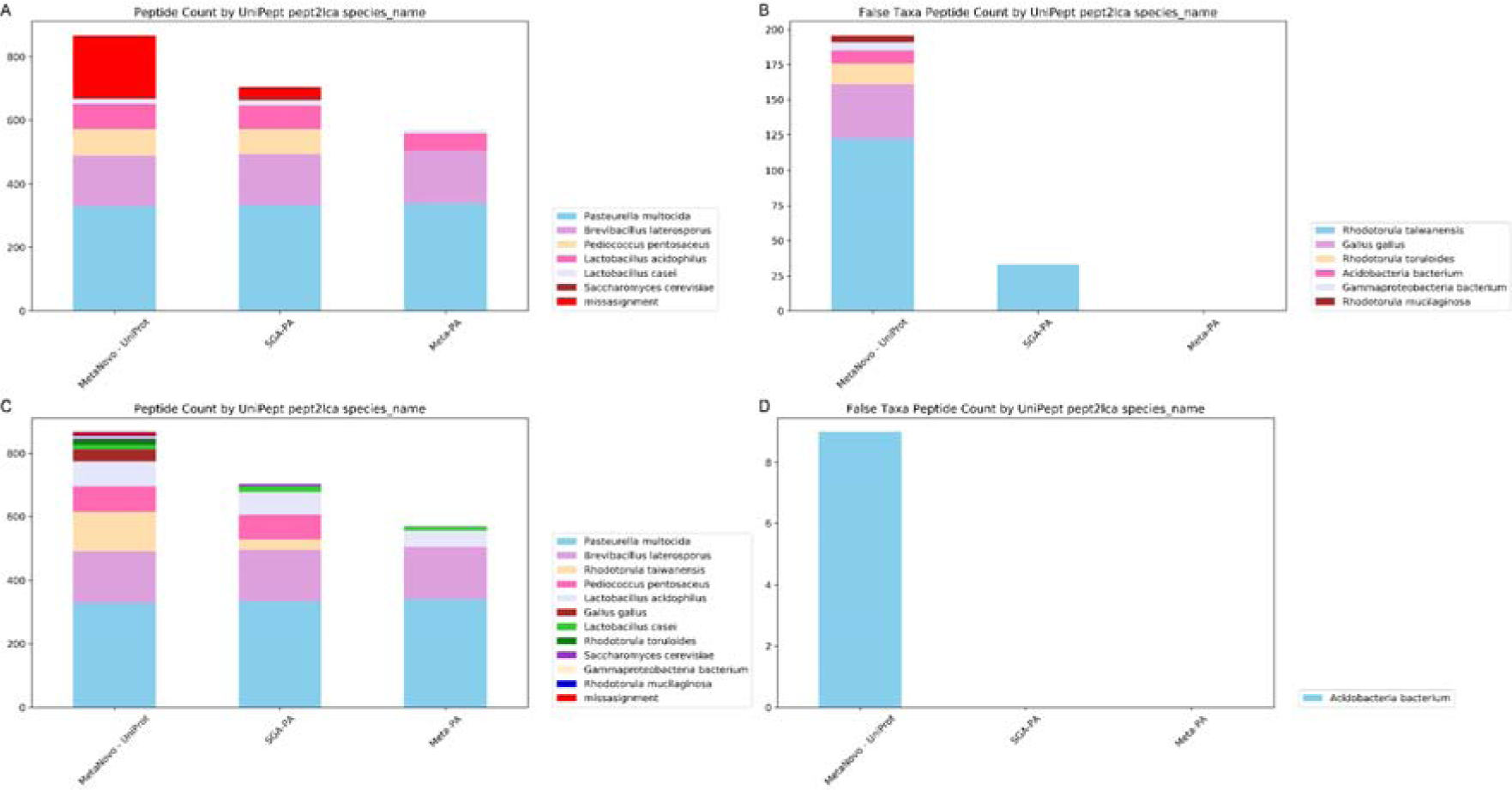
Percentages of misassigned peptides for all three runs. **A. MetaNovo** originally yielded a very high percentage of misassigned spectra at species level based on **UniPept** *pept2lca* analysis. **B.** Further analysis of the misassigned peptides indicated that most belonged to other *Rhodotorula* species. The *Gallus gallus* peptides were narrowed down to components of egg-yolk using **BLASTP** analysis. **C.** After allowing for other *Rhodotorula spp.*, *Gallus gallus* and *Gammaproteobacteria bacterium* peptides, a species-level misassignment rate of only 1.04 % was attained. **D.** 9 *Acidobacteria bacterium* peptides making up the final misassignment percentage of **MetaNovo** could be explained by the higher total number of identified peptides leading to more false hits at the same FDR, actual contaminants, or **UniPept** misassignments of natural variants in the data.

#### PEP Score analysis

Visual inspection of PEP score density plots shows a similar distribution of peptide PEP scores for peptides only identified in a specific run, and peptides that were identified in all runs, with the exception of peptides only identified using the **MetaPro-IQ/Metagenome** database, that showed a very broad distribution overlapping with the reverse hits (*See Supplementary Figure 1. Density plot of peptide PEP scores by group*).

Analysis of shared and exclusive peptide identifications in each group revealed significantly different distributions between 14049 peptides only identified using **MetaNovo** and reverse hits for the same run (p-value 6.33e-26), indicating true positive identifications in this set not found using the other approaches. 6525 peptides only identified using the matched metagenome did not have significantly different PEP scores from reverse hits (4.60e-01), suggesting a higher rate of false-positive hits. The difference between peptides exclusively identified with the **MetaPro-IQ/IGC** database and reverse hits was more significant (p-value 3.15e-20) but less so than for the **MetaNovo** comparison (*See Figure 4 D. Boxplot of Peptide PEP score distributions, Supplementary Table 2*).

#### Taxonomic comparisons

Peptide counts by lowest common ancestor were compared between different analysis runs using the output of UniPept *pept2lca*. After applying stringency criteria based on requiring at least 0.5 % of total classified peptides per included taxon, the highest number of characterised peptides by *taxon_name* was obtained using **MetaNovo/UniProt** (35169), followed by **MetaPro-IQ/Metagenome** (34768) and **MetaPro-IQ/IGC** (32325). A similar taxonomic distribution was obtained between runs (*See Figure 4 C).* Of the characterised peptides passing the stringency criteria, 4700 peptides belonged to *taxon_name* taxa that were identified exclusively by **MetaNovo**. These taxa are consistent with expected host peptides that were not identified by the other runs (no *Homo sapiens* peptides were identified by the other approaches). *See Supplementary Table 3. Peptide counts of taxa only identified with MetaNovo.* No taxa identified by the other runs were not also identified by **MetaNovo.** *See Supplementary Table 4. Peptide counts of taxa identified in all runs by UniPept pept2lca taxon_name.* When selecting only *Bacteria* “*superkingdom_name”* peptides lower numbers of peptides were identified by **MetaNovo** than for the other runs, with 27289, 30311 and 31591 annotated peptides for **MetaNovo**, **MetaPro-IQ/IGC**, and **MetaPro-IQ/Metagenome**, respectively. A similar taxonomic distribution was visualised for all three runs in this group. *See Supplementary Figure 2..* When aggregating peptide counts at the “*kingdom_name”* level, peptides from *Metazoa, Viridiplantae* and *Fungi* kingdoms were represented by all runs, but with much higher numbers represented by **MetaNovo.** These taxa are all known to play key roles in the human gut microbiome, as living or dietary components. *See Supplementary Table 5*. Similarly, aggregated to *Phylum* level, **MetaNovo** identified the same phyla as identified in either of the other runs, but with much higher numbers of *Chordata* peptides. Interestingly, both the **MetaNovo/UniProt** and **MetaProIQ/IGC** databases identified *Actinobacteria* - but these were not identified by the **MetaProIQ/Metagenome** database. *See Supplementary Table 6*.

### Analysis of a known microbial mixture using the MetaNovo software yields increased sensitivity compared to matched genomics databases

#### Protein and peptide identifications

The **MetaNovo** pipeline was run using the entire, unfiltered **UniProt** database (December 2019 release; ca. 180 million entries), resulting in the identification of 12529 target peptides and 2534 target protein groups, with 44.17 % of total MS/MS identified. These results are significantly better with 9.14 % more spectra assigned than the best performing genomic database reported by the original authors when run with the same **MaxQuant** settings (single predicted and annotated genomes assembly DB (SGA-PA)) that yielded 35.03 % identified MSMS, 10109 target peptides and 2099 target protein groups. The results were almost double those of the metagenome DB (Meta-PA) database that yielded 23.03 % identified MSMS, 6044 target peptides and 1087 target proteins (*See Supplementary **Figure 3******. A.***).

**Taxon identification rates.** Percentages of misassigned peptides were calculated for all three runs. **MetaNovo** originally yielded a very high percentage of misassigned spectra at species level based on **UniPept** *pept2lca* analysis. Further analysis of the characterised peptides indicated that most belonged to other *Rhodotorula* species, which corresponds to the authors’ original report of many non-canonical peptides identified in their samples using a 6-frame translated database, which could lead to **UniPept** assignment to related organisms. The identification of 38 *Gallus gallus* peptides was narrowed down to components of egg-yolk using **BLASTP** analysis (See *Supplementary Table 6*.). Personal correspondence with the authors confirmed that egg yolk was present in one of the growth media employed in the experiment. The *Pasteurella multicoda* strain analyzed in this study was a field isolate (author correspondence), and being a member of *Gammaproteobacteria*, natural sequence diversity in this strain might be compatible with a classification of *Gammaproteobacteria bacterium*. After allowing for other *Rhodotorula spp.*, *Gallus gallus* and *Gammaproteobacteria bacterium* peptides, a species-level misassignment rate of only 1.04 % was attained, with 0% error for all databases at genus and family level using the 0.5 % taxon-specific peptide stringency cutoff. The 9 *Acidobacteria bacterium* peptides making up the final misassignment percentage of **MetaNovo** could be explained by the higher total number of identified peptides leading to more false hits at the same FDR, actual contaminants, or **UniPept** misassignments of natural variants in the data.

#### Taxon assignment accuracy benchmarks based on mean-squared error show the highest accuracy for species-level assignments using MetaNovo

Mean Squared Error (MSE) scores were calculated for the relative proportion of MS/MS of each taxon to the total of each run compared to the expected proportion by CFU counting as a percentage. Scores closer to 0 indicate higher accuracy. MetaNovo had a poorer score than SGA-PA for genus level assignments, but the highest accuracy of the three runs for both family and species level assignments. MetaNovo outperformed the matched metagenomic database (Meta-PA) at all three taxonomic levels. After relabelling all *Rhodotorula* peptides as *Rhodotorula spp.* for the comparison, a further improvement in MSE scores was obtained, and corresponding to the author’s report of identifying many non-canonical peptides using the translated 6 frame database for the same organism, some of which may be homologous and identical to canonical peptides in other species in the same genus, and therefore were assigned to those organisms by UniPept. *See Table 1*.

**Table 1.** MetaNovo yields the highest accuracy for species-level annotations compared to matched genomic databases. Mean Squared Error (MSE) scores for the relative proportion of MSMS of each taxon to the total of each run compared to the expected proportion by CFU counting as a percentage. Scores closer to 0 indicate higher accuracy.

|  | Meta-PA | SGA-PA | MetaNovo - UniProt |
| --- | --- | --- | --- |
| **Family** | 398.89 | 143.58 | 106.08 |
| **Genus** | 261.64 | 78.44 | 87.32 |
| **Species** | 333.17 | 207.77 | 151.05 |
| **Species (relabelled)** | 333.17 | 198.79 | 138.03 |

#### Unexpected taxa BLAST analysis

Of the taxonomically assigned peptides across the three runs, 38 *Gallus gallus* proteins were the largest group of false assignments by the **MetaNovo** pipeline compared to the list of expected taxa. This group was chosen for protein **BLAST** analysis. For each peptide, BLAST results were filtered to only include exact matches (100 % identity). For each of these records, the organism names were identified, and 20 peptides that only matched to *Gallus gallus* were obtained as proteotypic peptides for this species, providing strong evidence at sequence level for this species in the data. 8 FASTA entries from the protein groups of the *Gallus gallus* peptides were obtained and their corresponding sequences were submitted for protein **BLAST**. For each query in the FASTA, the set of organisms for all matches with more than 90 % sequence identity was investigated - and 5 sequences matched only to *Gallus gallus.* These proteins include *Apovitellenin-1* (P02659), *Vitellogenin-1* (P87498, A0A1D5NUW2), *Vitellogenin-2* (P02845) and *Vitellogenin-3* (A0A3Q2U347). These are all known components of egg yolk.

#### Runtime performance measures of the MetaNovo pipeline on the 9MM samples

On a 32 core Intel Core Processor (Broadwell, IBRS) server with 236Gi and 32 threads set in the **MetaNovo** config for the analysis of the two 9MM raw files described above against the December 2019 **UniProt** release (ca 180 million sequences) a total runtime of 0:09:50:7.2816 was recorded.

## Discussion

On re-analysis of a set of Human Mucosal-luminal interface samples, comparative identification rates were achieved, with a >2 % increase in the number of MS/MS at the same FDR - but now simultaneously identifying multiple human origin peptides missed by the previous approaches - while not leaving out taxa identified by the other runs. The large number of exclusive peptides only identified by *MetaNovo*, can be explained by the previous approaches not including host sequences in the database, with potential spurious assignments of host peptides to microbial database entries for those runs - a known risk for databases that are not comprehensive (recall the *Apis mellifera* reanalysis example discussed previously) [11]. These spurious PSMs would have a higher PEP score distribution, overlapping with the reverse hit database (as all are spurious sequences) - thus leading to a decreased rate of target peptide identifications at the given FDR cutoff. This can explain the lower MS/MS % identification rate of the previous approaches. The improved error distribution of exclusive peptides identified using **MetaNovo**, and the error distribution of exclusive peptides that were identified with the metagenomics database not being significantly different from the distribution of reverse hits for the same run, support this view. As **MetaNovo** did not identify any exclusive taxa other than host compared to the other runs, but with higher numbers of viridiplantae and fungi taxa, it points to improved characterization of non-bacterial components while maintaining an accurate characterization of the bacterial component without over-estimating taxonomic diversity. These other components of the gut microbiome may yield key insights into complex inter-specific interactions that may not be evident with approaches that target only limited taxonomies. Lower numbers of Bacterial peptide identifications by **MetaNovo** compared to the other two approaches are likely due to the fact that both previous databases are more comprehensive at the Bacterial level, while the **MetaNovo** algorithm aims to be globally representative, and increase the percentage of assigned spectra overall. Thus, future updates to the algorithm will aim to increase the comprehensiveness of the generated database so that the output can compete with taxonomically focused databases and further increase the global sensitivity of assignments.

Taxonomic assignment of peptides to other *Rhodotorula* species than the expected strain by **UniPept**, for both **MetaNovo** (143 peptides) and **SGA-PA** (33 peptides), indicate the limitations of the **UniPept** approach that is based on the transfer of reference annotations to the natural sequence diversity of field isolates. The higher number of *Rhodotorula* peptides are consistent with the lower sequence coverage reported for the *R. glutinis* strain (x12) used for the **SGA-PA** database. It is notable that the **MetaNovo** approach was able to yield improved performance over full genome sequencing by leveraging sequence information of related strains without prior expectation.

The identification of proteotypic peptides for *Gallus gallus*, as well as the identification of 5 proteotypic protein groups all related to egg yolk, was strong evidence for egg yolk contamination in the 9MM samples. To confirm this, personal correspondence with the authors confirmed the presence of egg yolk in one of the growing media. Thus, the identification of putative contaminants that may have otherwise been missed during database creation, confirms the usefulness of the approach for cases where rare and unknown contaminants would otherwise be missed.

Higher rates of MS/MS assignments yielded by the **MetaNovo** approach compared to the matched genomic approaches (both **MetaPro-IQ/Metagenome** and **SGA-PA**) indicate the power of the **MetaNovo** algorithm to leverage publicly available taxonomic and sequence data that may be vastly more representative of expressed proteomes and natural sequence diversity, than omics approaches that do not directly reflect the actual sequences present at the proteome level. Leveraging the high numbers of curated sequences available, while not replacing matched genomics, can certainly be considered a viable alternative where sequencing is not available, and where novel sequence polymorphism is of minor concern. Further advantages of a protein-only database creation approach include the disadvantage of 16S sequencing approaches that only focus on the bacterial component or any genomics approach that would not take into account secretory proteins that are generated in other biological compartments, as well as dietary components that may not have corresponding genetic material present in the microbiome but play an important role in gut homeostasis as well as being present at the protein level. By producing more representative but still focused databases, the spurious assignment of MS/MS due to database incompleteness will be reduced, leading to an increase in database search sensitivity and accuracy.

**MetaNovo** yielded higher accuracy of taxonomic assignment in terms of relative proportions of species in the samples illustrated with the MSE analysis relative to expected proportions based on CFU counts, with comparable error rates and a vast (almost double) increase in assigned spectra with this approach. While the reliance on annotated sequences will miss many novel sequence polymorphisms that may only be identified by matched genome sequencing, the improvement in search sensitivity with the approach is promising.

It can be argued that the use of **UniProt** (**SwissProt** and **trEMBL**), although comprehensive, is biased in database composition when compared to much larger databases such as **NCBI nr** which contain many more proteins and species from clinical and environmental sources. Further work must be done to tailor the **MetaNovo** algorithm to work with non-**UniProt** FASTA databases and to ensure that it is performant with such bigger databases, as well as the analysis of metagenomic databases that may benefit from a two-step search approach.

## Conclusion

**MetaNovo** makes characterising complex mass spectrometry data with an extremely large, space feasible, allowing for novel peptide identifications, even when genomic information is not available, and improving the accuracy and affordability of current clinical proteomic and metaproteomic analysis methods by leveraging the rapidly growing number of available sequences from genome and metagenome sequencing efforts. By tailoring a database for the samples to be analysed from **UniProt** in an unbiased manner based on a direct estimate of proteins and taxa present in the data, **MetaNovo** allows proteomic identification to be carried out, unlimited by prior expectation, with comparable results to matched metagenomic databases.

We have shown that using **MetaNovo,** a sensitive characterization of the human gut metaproteome allows for a simultaneous snapshot of the proteins at play in both the host and the microbiome. Finally, the ability to search an extremely large database such as **UniProt** provides the potential to identify microbiota and biomarkers that may be missed using limited or hand-selected databases. **MetaNovo** results on samples of known mixture compare favourably to matched genomics approaches, increasing the sensitivity without overestimating taxa present in the samples, and simultaneously identifying unknown contaminants that may lead to downstream errors if not identified.

By estimating taxonomic and peptide level information on microbiome samples directly from tandem mass spectrometry data, in combination with the entire **UniProt** database, **MetaNovo** enables simultaneous identification of human, bacterial, fungal, and other eukaryotic proteins in a sample, thus potentially allowing clinically-important correlations to be drawn between changes in microbial protein abundance and change in the host proteome, within a single analysis.

Using the freely-available **MetaNovo** algorithm, researchers will be able to focus on downstream analysis to extract biological meaning at the level of expressed proteomes in microbiome samples, whilst minimising reliance on prior knowledge, or external information such as whole-genome sequencing. It is hoped that the addition of probabilistic protein inference after the first search prior to the second search by the **MetaNovo** approach, will extend the state of the art of two-step iterative database searches in mass-spectrometry metaproteomics, and lead to future innovations in the field.

## Availability and requirements

Project name: MetaNovo

Project home page: https://github.com/uct-cbio/proteomics-pipelines

Operating system: Linux

Programming language: Bash and Python

Other requirements: Docker/Singularity/PBS

Licence: MIT

## List of abbreviations

NSAF: Normalised Spectral Abundance Factor
SAF: Spectral Abundance Factor
MLI: Mucosal-luminal interface
TIC: total ion chromatogram
QE: Q-Exactive Orbitrap mass spectrometer
insol: in solution
FASP: filter-assisted sample processing
PEP: Posterior error probability

## Declarations

### Ethics statement

The current publication relies on re-analysis of publicly available mass-sepctrometry data.

### Consent for publication

Not applicable.

### Availability of data and materials

MetaNovo software is available from GitHub^3^ and can be run as a standalone Singularity or Docker container available from the Docker Hub^4^. The mass spectrometry proteomics data used to benchmark the software have been deposited to the ProteomeXchange Consortium via the PRIDE [24] partner repository with the dataset identifier **PXD030708**.

### Funding and acknowledgements

MGP would like to thank the NRF for an MSc grant (NRF BFG 93665) and Professor Nicola Mulder for a PhD grant. DLT was supported by the

South African Tuberculosis Bioinformatics Initiative (SATBBI), a Strategic Health Innovation Partnership grant from the South African Medical Research Council and South African Department of Science and Technology. JMB thanks the NRF for a South African Research Chair grant.

### Author contributions

MGP - design of pipeline; Bioinformatic data analysis; wrote manuscript AJMN – Bioinformatic data analysis; experimental data generation SF - Experimental data generation SG - Experimental data generation JW - Experimental data generation

DT - DirecTag configuration; Taxonomic weighting strategy

NM - Design of overall study; design of pipeline; wrote manuscript; corresponding author JB – Design of overall study; design of pipeline; wrote manuscript; corresponding author

## Footnotes

1 https://github.com/uct-cbio/proteomics-pipelines

2 https://hub.docker.com/r/thyscbio/metanovo

3 https://github.com/uct-cbio/proteomics-pipelines

4 https://hub.docker.com/r/thyscbio/metanovo

