## Supplemental Data for "*MetaNovo:* a probabilistic approach to peptide discovery in complex metaproteomic datasets"

**MetaNovo Supplementary Data**

**1. Supplementary Tables**

|  |  | **SwissProt** | **TREMBL** | **Total** | **%** |
| --- | --- | --- | --- | --- | --- |
| **Archaea** |  | 19581 | 3933054 | 3952635 | 2.20 |
| **Bacteria** |  | 334357 | 128577841 | 128912198 | 71.69 |
| **Viruses** |  | 16988 | 4303549 | 4320537 | 2.40 |
| **Other** |  | 0 | 1687026 | 1687026 | 0.94 |
| **Eukaryota** | **Human** | 20368 | 168078 | 188446 | 0.10 |
|  | **Other Mammalia** | 46839 | 3179841 | 3226680 | 1.79 |
|  | **Other Vertebrata** | 18434 | 4807649 | 4826083 | 2.68 |
|  | **Viridiplantae** | 40216 | 9314135 | 9354351 | 5.20 |
|  | **Fungi** | 34262 | 11564971 | 11599233 | 6.45 |
|  | **Insecta** | 9354 | 3811058 | 3820412 | 2.12 |
|  | **Nematoda** | 5015 | 1847926 | 1852941 | 1.03 |
|  | **Other** | 16154 | 6055433 | 6071587 | 3.38 |
| **Total** |  | 561568 | 179250561 | 179812129 | 100.00 |

**Supplementary Table 1. Taxonomic distribution by Kingdom of the UniProt database.** The 2019_11 release of **UniProt** consists of ca. 180 million protein sequences, the majority of which are Bacteria.

|  | **metanovo - UniProt: Shared** | **metanovo - UniProt: Only** | **metanovo - UniProt: Reverse** | **metaproiq - igc: Shared** | **metaproiq - igc: Only** | **metaproiq - igc: Reverse** | **metaproiq - metagenome: Shared** | **metaproiq - metagenome: Only** | **metaproiq - metagenome: Reverse** |
| --- | --- | --- | --- | --- | --- | --- | --- | --- | --- |
| **metanovo - UniProt: Shared** | -1.000000e+00 | 3.734901e-87 | 1.729789e-39 | 3.773263e-32 | 4.537713e-216 | 1.541020e-60 | 7.627574e-72 | 0.000000e+00 | 1.569875e-53 |
| **metanovo - UniProt: Only** | 3.734901e-87 | -1.000000e+00 | 6.330437e-26 | 1.158218e-32 | 3.620188e-64 | 1.281311e-41 | 2.606551e-222 | 0.000000e+00 | 1.481513e-36 |
| **metanovo - UniProt: Reverse** | 1.729789e-39 | 6.330437e-26 | -1.000000e+00 | 7.918533e-34 | 3.192363e-11 | 2.014311e-01 | 5.813792e-49 | 3.362623e-02 | 3.147086e-01 |
| **metaproiq - igc: Shared** | 3.773263e-32 | 1.158218e-32 | 7.918533e-34 | -1.000000e+00 | 9.163935e-153 | 9.235974e-53 | 1.125309e-194 | 0.000000e+00 | 1.536492e-46 |
| **metaproiq - igc: Only** | 4.537713e-216 | 3.620188e-64 | 3.192363e-11 | 9.163935e-153 | -1.000000e+00 | 3.148180e-20 | 0.000000e+00 | 4.475586e-267 | 1.827976e-17 |
| **metaproiq - igc: Reverse** | 1.541020e-60 | 1.281311e-41 | 2.014311e-01 | 9.235974e-53 | 3.148180e-20 | -1.000000e+00 | 2.271949e-73 | 6.870360e-01 | 7.870152e-01 |
| **metaproiq - metagenome: Shared** | 7.627574e-72 | 2.606551e-222 | 5.813792e-49 | 1.125309e-194 | 0.000000e+00 | 2.271949e-73 | -1.000000e+00 | 0.000000e+00 | 3.725671e-65 |
| **metaproiq - metagenome: Only** | 0.000000e+00 | 0.000000e+00 | 3.362623e-02 | 0.000000e+00 | 4.475586e-267 | 6.870360e-01 | 0.000000e+00 | -1.000000e+00 | 4.598942e-01 |
| **metaproiq - metagenome: Reverse** | 1.569875e-53 | 1.481513e-36 | 3.147086e-01 | 1.536492e-46 | 1.827976e-17 | 7.870152e-01 | 3.725671e-65 | 4.598942e-01 | -1.000000e+00 |

**Supplementary Table 2. Peptide PEP Score comparison between shared, exclusive and reverse hits of different approaches.** Peptides identified only using **MetaNovo** had a significantly different PEP score distribution by p-value compared to reverse hits for the same run after applying Dunn’s test post-hoc analysis after Kruskal-Wallis analysis for variance. Kruskal-Wallis test indicated significant differences between groups (

KruskalResult statistic=10883.939894469078, p value=0.0).

|  | **metanovo - UniProt** | **metaproiq - igc** | **metaproiq - metagenome** |
| --- | --- | --- | --- |
| **taxon_name** |  |  |  |
| ***Mammalia*** | 1446 | 0 | 0 |
| ***Craniata*** | 563 | 0 | 0 |
| ***Sarcopterygii*** | 457 | 0 | 0 |
| ***Catarrhini*** | 399 | 0 | 0 |
| ***Homo sapiens*** | 395 | 0 | 0 |
| ***Metazoa*** | 378 | 0 | 0 |
| ***Simiiformes*** | 318 | 0 | 0 |
| ***Homininae*** | 280 | 0 | 0 |
| ***Eukaryota*** | 254 | 0 | 0 |
| ***Euarchontoglires*** | 210 | 0 | 0 |
| **Total** | 4700 | 0 | 0 |

**Supplementary Table 3. Peptide counts of taxa only identified with MetaNovo.** After applying stringency criteria based on requiring at least 0.5 % of all classified peptides per taxon, 4700 peptides belonged to taxa identified exclusively to **MetaNovo**. These peptides of *Chordata* origin are consistent with host peptides that were not identified by the other runs (no *Homo sapiens* peptides were identified by the other approaches). Including the correct taxa in the database, including host sequences, are important to ensure that sequences are not misassigned due to database non-representativity influencing the selection of top PSMs per MS/MS during target-decoy searches. No taxa identified in the other runs were not identified by **MetaNovo**.

|  | **metanovo - UniProt** | **metaproiq - igc** | **metaproiq - metagenome** |
| --- | --- | --- | --- |
| **taxon_name** |  |  |  |
| ***Bacteroides*** | 6025 | 6531 | 7004 |
| ***Clostridiales*** | 5793 | 6698 | 6602 |
| ***Bacteria*** | 4595 | 4838 | 5163 |
| ***Bacteroidales*** | 3293 | 3681 | 3909 |
| ***root*** | 3180 | 2014 | 3177 |
| ***Mammalia*** | 1446 | 0 | 0 |
| ***Faecalibacterium prausnitzii*** | 1235 | 1404 | 1383 |
| ***Bacteroides plebeius*** | 711 | 794 | 873 |
| ***Bacteroides fragilis*** | 715 | 778 | 866 |
| ***Bacteroidetes*** | 770 | 799 | 866 |
| ***Firmicutes*** | 706 | 805 | 732 |
| ***Clostridium*** | 580 | 733 | 717 |
| ***Craniata*** | 563 | 0 | 0 |
| ***Alistipes putredinis*** | 419 | 479 | 532 |
| ***Alistipes*** | 402 | 457 | 483 |
| ***Ruminococcaceae*** | 366 | 464 | 470 |
| ***Sarcopterygii*** | 457 | 0 | 0 |
| ***Lachnospiraceae*** | 364 | 423 | 430 |
| ***Erysipelotrichaceae*** | 367 | 384 | 407 |
| ***Catarrhini*** | 399 | 0 | 0 |
| ***Homo sapiens*** | 395 | 0 | 0 |
| ***Metazoa*** | 378 | 0 | 0 |
| ***Bacteroides stercoris*** | 247 | 276 | 329 |
| ***Simiiformes*** | 318 | 0 | 0 |
| ***Bacteroides uniformis*** | 267 | 286 | 313 |
| ***Homininae*** | 280 | 0 | 0 |
| ***Bacteroides caccae*** | 216 | 243 | 270 |
| ***Eukaryota*** | 254 | 0 | 0 |
| ***Bacteroidia*** | 218 | 238 | 242 |
| ***Euarchontoglires*** | 210 | 0 | 0 |
| **Total** | 35169 | 32325 | 34768 |

Supplementary Table 4. Peptide counts of taxa identified in all runs by UniPept pept2lca taxon_name**.** No peptides identified by the other runs were not identified by **MetaNovo.**

|  | **metanovo - UniProt** | **metaproiq - igc** | **metaproiq - metagenome** |
| --- | --- | --- | --- |
| **kingdom_name** |  |  |  |
| **Metazoa** | 5063 | 178 | 187 |
| **Viridiplantae** | 119 | 2 | 1 |
| **Fungi** | 35 | 5 | 3 |

Supplementary Table 5. Peptide counts of taxa by UniPept pept2lca kingdom_name. When aggregating peptide counts at *kingdom_name* level, peptides from *Metazoa, Viridiplantae* and *Fungi* kingdoms were represented by all runs, but with much higher identification rates by **MetaNovo.** These taxa are all known to play key roles in the human gut microbiome, as living or dietary components.

|  | **metanovo - UniProt** | **metaproiq - igc** | **metaproiq - metagenome** |
| --- | --- | --- | --- |
| **phylum_name** |  |  |  |
| ***Bacteroidetes*** | 14750 | 16359 | 17603 |
| ***Firmicutes*** | 10787 | 12494 | 12033 |
| ***Chordata*** | 4645 | 170 | 180 |
| ***Proteobacteria*** | 405 | 407 | 399 |
| ***Actinobacteria*** | 265 | 328 | 0 |
| **Total** | 30852 | 29758 | 30215 |

Supplementary Table 6. Peptide counts of taxa by UniPept pept2lca phylum_name. **MetaNovo** identified the same phyla as identified in either of the other runs, but with a much higher count of Chordata peptides. Interestingly, both the **MetaNovo/UniProt** and **MetaProIQ/IGC** databases identified *Actinobacteria* - but these were not identified by the **MetaProIQ/Metagenome** database.

|  | **MetaNovo** | **SGA-PA** | **Meta-PA** |
| --- | --- | --- | --- |
| **species_name** |  |  |  |
| ***Rhodotorula taiwanensis*** | 123 | 33 | 0 |
| ***Gallus gallus*** | 38 | 0 | 0 |
| ***Rhodotorula toruloides*** | 15 | 0 | 0 |
| ***Acidobacteria bacterium*** | 9 | 0 | 0 |
| ***Gammaproteobacteria bacterium*** | 6 | 0 | 0 |
| ***Rhodotorula mucilaginosa*** | 5 | 0 | 0 |
| **Total** | 196 | 33 | 0 |

**Supplementary Table 6.** False Taxa Peptide Count by UniPept pept2cla species_name

**2. Supplementary Figures**

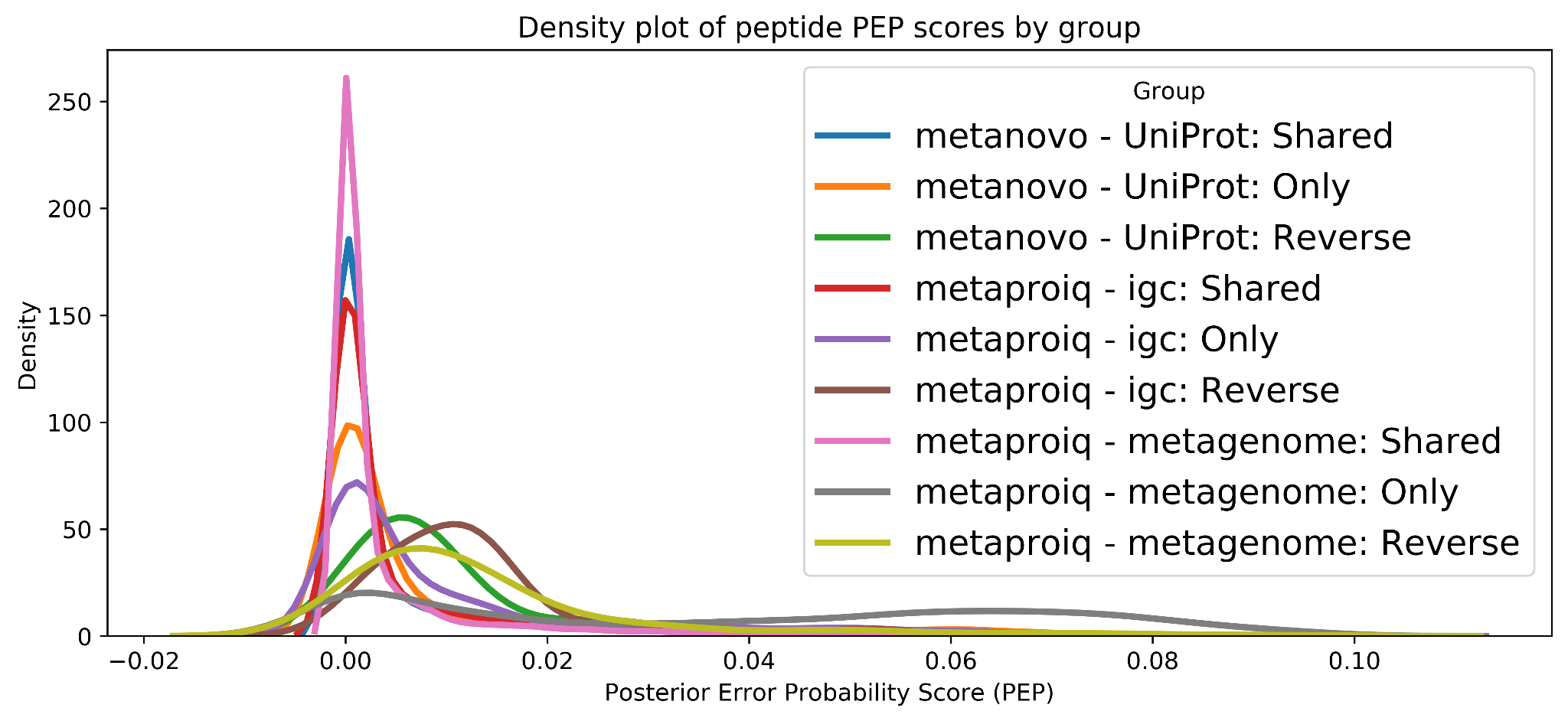

Supplementary Figure 1. Density plot of peptide PEP scores by group

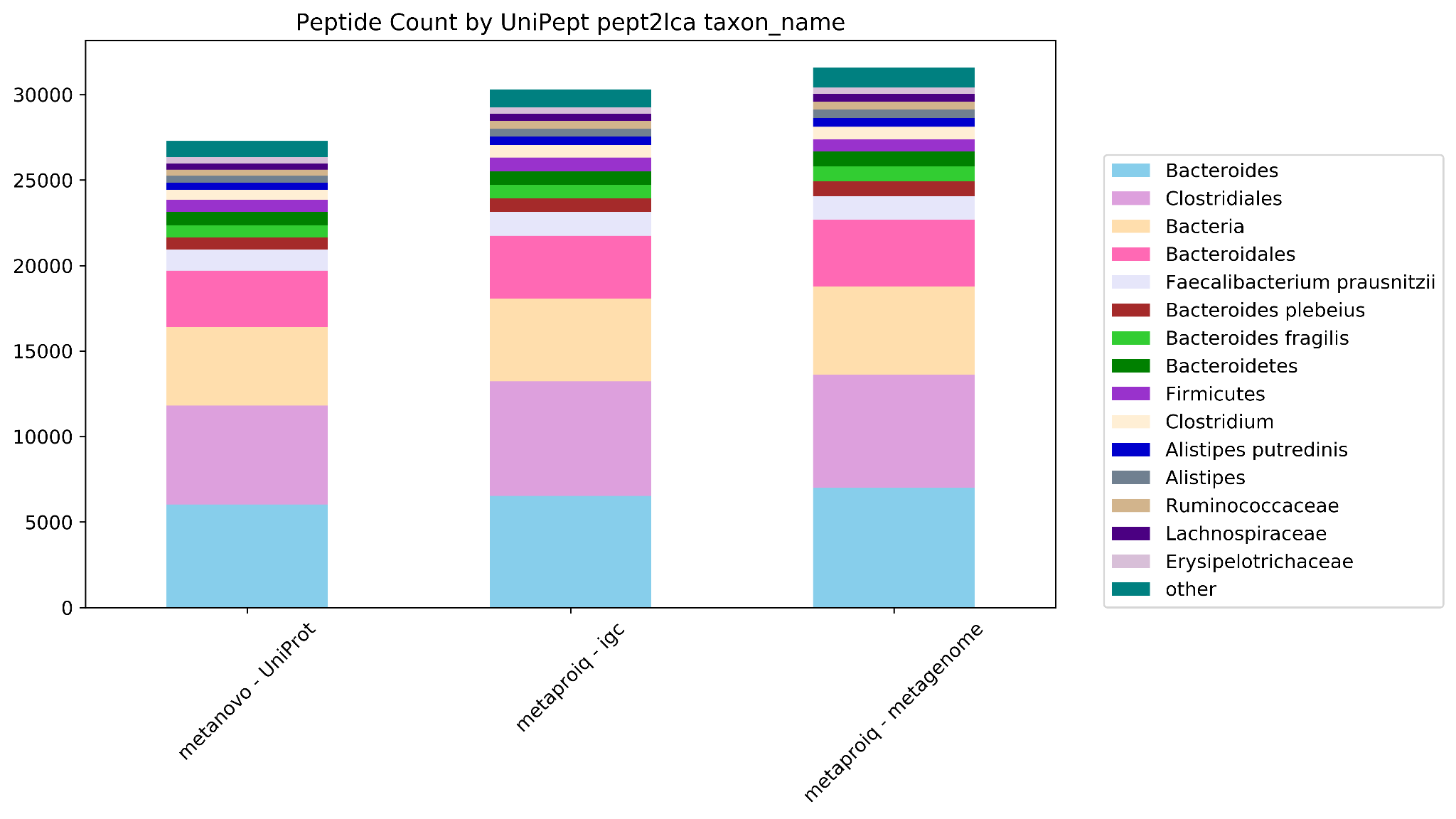

Supplementary Figure 2. Peptide counts by UniPept *pept2lca* taxon_name for all runs, *Bacteria* only

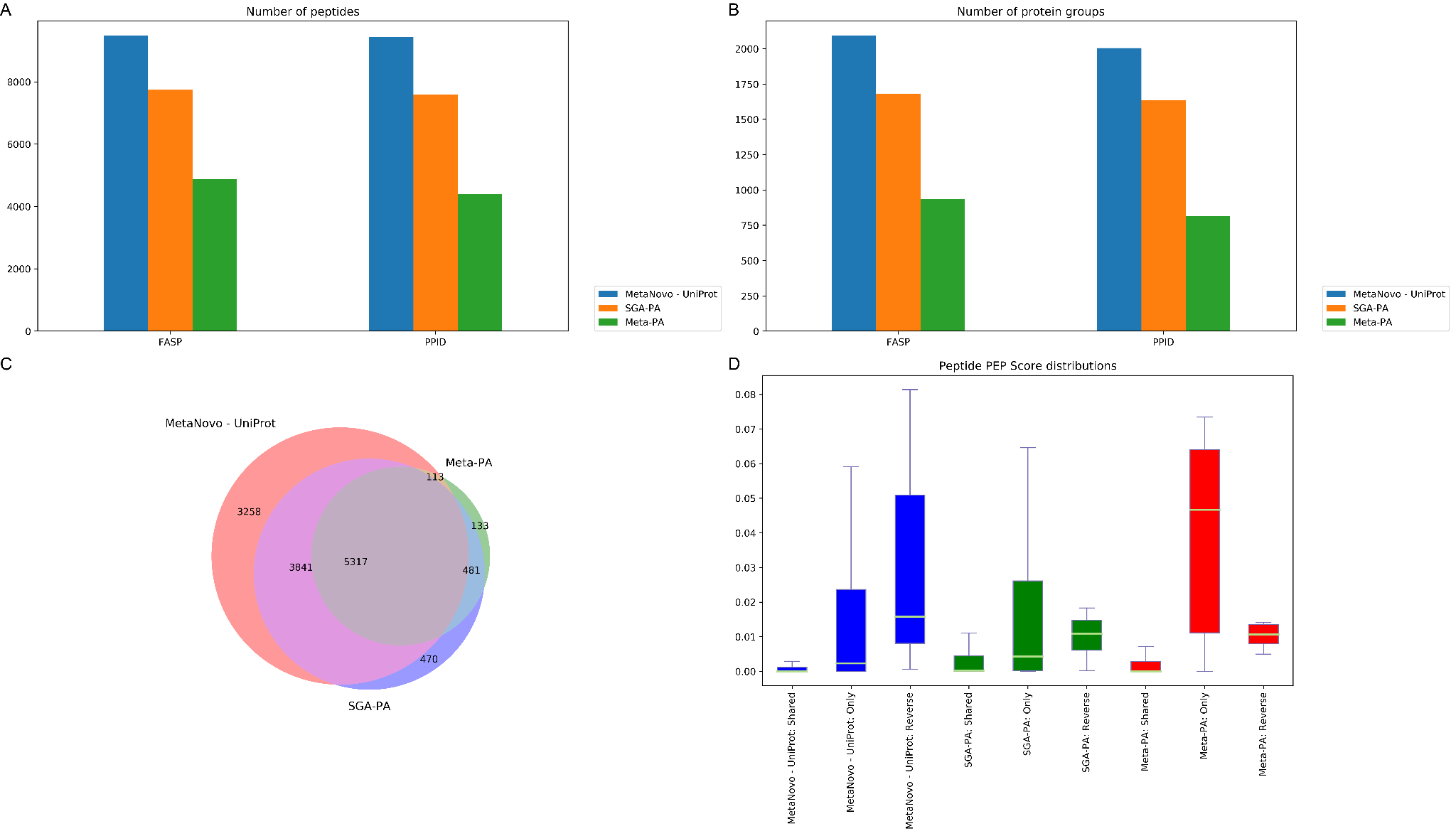

**Supplementary Figure 3. A.** Bar chart of peptide identifications. Many more peptides were obtained with **MetaNovo**. **B.** Bar chart of protein group identifications, showing a similar pattern to peptide hits. **C.** Venn diagram of peptide identifications across the three runs shows significant overlap between all three runs, and a very small percentage of hits identified by the other runs that were not identified by **MetaNovo** as well.. **D.** Peptides exclusively identified using the **MetaNovo** database had a lower median PEP score and better distribution than the exclusively identified peptides in the other two runs. The boxes extend from the lower to the upper quartile, and the whiskers represent 1.5 times the interquartile range (IQR) below and above the first and third quartiles, respectively.
